# RamaNet: Computational *de novo* helical protein backbone design using a long short-term memory generative neural network

**DOI:** 10.1101/671552

**Authors:** Sari Sabban, Mikhail Markovsky

**Affiliations:** Department of Biological Sciences/Faculty of Science, King Abdulaziz University, Jeddah, Makka, Kingdom of Saudi Arabia; Electrical and Computer Engineering Department/College of Engineering, Effat University, Jeddah, Makka, Kingdom of Saudi Arabia; North Caucasian Federal Scientific Center of Horticulture, Viticulture, Wine-making, Krasnodar, Russian Federation

**Keywords:** *De novo* protein design, Neural Networks, Long Short-Term Memory Neural Network, Deep Learning, Helical Proteins, Protein backbone

## Abstract

The ability to perform *de novo* protein design will allow researchers to expand the variety of available proteins. By designing synthetic structures computationally, they can utilise more structures than those available in the Protein Data Bank, design structures that are not found in nature, or direct the design of proteins to acquire a specific desired structure. While some researchers attempt to design proteins from first physical and thermodynamic principals, we decided to attempt to test whether it is possible to perform *de novo* helical protein design of just the backbone statistically using machine learning by building a model that uses a long short-term memory (LSTM) architecture. The LSTM model used only the *ϕ* and *ψ* angles of each residue from an augmented dataset of only helical protein structures. Though the network’s generated backbone structures were not perfect, they were idealised and evaluated post generation where the non-ideal structures were filtered out and the adequate structures kept. The results were successful in developing a logical, rigid, compact, helical protein backbone topology. This paper is a proof of concept that shows it is possible to generate a novel helical backbone topology using an LSTM neural network architecture using only the *ϕ* and *ψ* angles as features. The next step is to attempt to use these backbone topologies and sequence design them to form complete protein structures.

**Author summary:** This research project stemmed from the desire to expand the pool of protein structures that can be used as scaffolds in computational vaccine development, since the number of structures available from the Protein Data Bank was not sufficient to allow for great diversity and increase the probability of grafting a target motif onto a protein scaffold. Since a protein structure’s backbone can be defined by the *ϕ* and *ψ* angles of each amino acid in the polypeptide and can effectively translate a protein’s 3D structure into a table of numbers, and since protein structures are not random, this numerical representation of protein structures can be used to train a neural network to mathematically generalise what a protein structure is, and therefore generate new a protein backbone. Instead of using all proteins in the Protein Data Bank a curated dataset was used encompassing protein structures with specific characteristics that will, theoretically, allow them to be evaluated computationally. This paper details how a trained neural network was able to successfully generate helical protein backbones.

## Introduction

The concept of amino acid sequences folding into globular protein molecules allows for proteins’ large functional diversity, mediating all the functional aspects of living organisms, thus winning themselves attention from biochemists for decades. The fusion of machine learning and deep learning with computational biology is accelerating research in both fields and bringing humanity closer to the setup of performing most biological research quickly, cheaply, and safely *in silico*, while only translating the very crucial aspects of it. Having access to a large database of protein crystal structures has led to the use of machine learning to design proteins computationally.

*De novo* protein design (i.e. from the beginning) is very well explained in this review [9]. Proteins fold into a specific shape depending on the sequence of their amino acids, and of course shape dictates function. The driving forces that allow proteins to fold are the hydrogen bond interactions within the backbone and between the side chains, the Van der Waals forces, and principally the interaction of hydrophobic side chains within the core. The space of all possible sequences for all protein sizes is extremely large (as an example there are 20^200^ possibilities for a 200-residue protein). Thus, is it not surprising that natural proteins exist in clusters close to each other, which is logical since proteins would evolve away from a central functional protein to fold correctly and acquire new folds and functions, rather than go through the tedious ordeal of finding a totally new protein structure within the space of all possibilities. Thus, even though the Protein Data Bank adds about 10,000 new structures to its repository every year, most of these new structures are not unique folds.

The relationship between the sequence of a protein and its specific structure is understood, but we still lack a unified absolute solution to calculate one from the other. Hence why some research groups generated man-made protein designs by altering already existing natural proteins [5], since randomly finding a functionally folded protein from the space of all possible protein sequences is more or less statistically impossible. On the other hand, other researchers have attempted *de novo* protein design by designing a topology from assembling short sequence peptide fragments taken from natural protein crystal structures [14] [8], these fragments are calculated statistically depending on the secondary structures they are found in. Sometimes this fragment system is combined with first physical principals to model the loops between secondary structures to achieve a desired three-dimensional topology [13]. Others have used parametric equations to study and specify the desired protein geometry [16] [7] [6] [10] [11] [17]. These solutions employ an energy function, such as REF2015, that uses some fundamental physical theories, statistical mechanical models, and observations of protein structures to approximate the potential energy of a protein [1]. Knowing the protein potential energy allows us to guide our search for the structure of a protein given its sequence (the structure resides at the global energy minima of that protein sequence) thus attempting to connect the sequence of a protein with its structure. The critical tool is the energy function, the higher its accuracy the higher our confidence in knowing the computed structure is the real natural structure. Thus, using the energy function to perform structure prediction (going from a known sequence to find the unknown three-dimensional structure) can also be used to perform fixed-backbone design (going from a known three-dimensional structure to find the sequence that folds it). This is where this paper comes in. Where as in *de novo* design neither backbone nor sequence is known, knowing one results in finding the other using the same energy function [9], and a good starting point is to design the backbone.

Other researchers have used machine learning for protein sequence design, employing the constraints (C*α*-C*α* distances) as the input features for the network and using a sliding window to read a sequence of residues, getting their types and constraints then predicting the next one giving the output prediction as an amino acid sequence [20], this architecture reported an accuracy of 38.3% and performs what is called sequence design: designing a sequence for a backbone, so when the protein is synthesised it folds to that backbone. In fact, in the [13] paper the protocol first generated an all-valine backbone, then sequence designed that backbone. In this paper, we want to computationally generate a backbone so it can be sequence designed using other protocols such as RosettaDesign[24] [25] or the protocol from [20].

The protein’s backbone can be folded using the *ϕ* and *ψ* angles, which are the angles between C*α* and N for *ϕ* and C*α* and C for *ψ* that primarily move the amino acid’s backbone (not the side chains), and thus can be used as features to control the topology of a backbone. They were even one of the features used to fold proteins and predict their structures in AlphaFold [23].

But the question is: how do we decide the ideal angles for a helix, the length of each helix, the number of helices, as well as the lengths and angles of the loops between the helices that will also result is a compact folded protein backbone. These numerous values can be solved statistically using neural networks. Especially that we want to use the structures in the PDB to forward design (rather then discover) new protein folds that are not a result of evolution.

The deep neural network architecture that we chose was a long short-term memory (LSTM) network [15]. The LSTM is usually used in natural language and data sequence processing, but in our model the LSTM was used to generate a sequence of *ϕ* and *ψ* angles. The model was constructed as a stack of LSTM layers, followed by fully connected layers, followed by a mixture density network (MDN) [31] and worked by using random noise numbers (a latent) as input to build the values for the *ϕ* and *ψ* angles [2].

Our effort in this paper was to use deep learning to learn the general fold of natural proteins, and using this generalising statistical concept to design novel protein backbone topologies, thus only getting the three dimensional backbone structure so it can be sequence designed using other protocols. Our research at this moment was a proof of concept and only concerned with getting a new and unique folded ideal helical protein backbone rather than a protein with a specific sequence, nor a function, nor a specific predetermined structure. Our system resulted in random yet compact helical backbone topologies only.

## Methods

The following steps were used to generate the augmented training dataset, along with details of the neural network architecture and how the output was optimised then evaluated.

### Data generation

The entire PDB database was downloaded on 28^th^ June 2018 (∼150,000 structures), and each entry was divided into its constituent chains resulting in individual separate structures (i.e: each PDB file had only a single chain). Each structure was analysed and chosen only if it contained the following criteria: contained only polypeptides, had a size between 80 and 150 amino acids without any breaks in the chain (a continuous polypeptide), a sum of residues that made up helices and sheets were larger than the sum of amino acids that made up loops, and the final structure having an Rg (radius of gyration) value of less than 15 88 Å. The chosen structures were then further filtered by a human to ensure only the desired structure concepts were selected, removing the structures that slipped through the initial computational filter. Furthermore, a diversity of structure folds were achieved rather than numerous repeats of the same fold (the haemoglobin fold was quite abundant). In previous attempts, a mixture of different structure classes were used, where some structures were only helices, some were only sheets, and the remaining were a mix of the two. However, that proved challenging in optimising the network, and as such a dataset made up of only helical protein structures was chosen for this initial proof of concept. The final dataset had 607 ideal helical structures. These structures were then cleaned (non-amino acid atoms were removed) in preparation to push them through the Rosetta version 3 modelling software that only takes in polypeptide molecules.

### Data augmentation

These 607 structures were augmented using the Rosetta FastRelax protocol [19]. This protocol performs multiple cycles of packing and minimisation. In other words, the protocol performs small slight random angle moves on the backbone and side chains in an attempt to find the lowest-scoring variant, but the random backbone angle moves is what we were after. Its originally intended function was to move a structure slightly to find the conformation of the backbone and side chains that corresponds to the lowest energy state as per the REF15 energy function. Since the protocol performs random moves, a structure relaxed on two separate occasions will result in two molecules that look very similar with similar minimum energy scores, but technically have different *ϕ* and *ψ* angle values. This is the concept we used to augment our structures, and each structure was relaxed 500 times to give a final dataset size of 303,500 structures.

### Feature extraction

Using only the *ϕ* and *ψ* angle values from a crystal structure it was possible to re-fold a structure back to its correct native fold, thus these angles were the only relevant features required to correctly fold a structure, Figure 1A details the range of angles in the un-augmented data. Each amino acid’s *ϕ* and *ψ* angle values were extracted and tabulates as in Table 1. This was the dataset used to train the neural network.

**Table 1.**
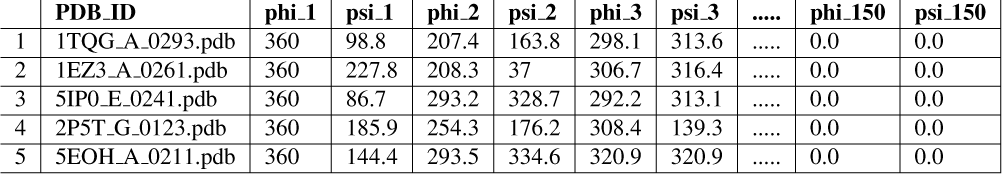
The PS Helix 500.csv dataset: The first five examples of the PS Helix 500.csv dataset showing the PDB ID chain augmentation number, residue 1 *ϕ* angle, residue 1 *ψ* angle, all the way to residue 150. 360.0° was used for the first missing angle while 0.0° was used to represent no residues.

**Table 2.** Structure packing scores: This table summarises the packing score of each structure, calculated as the average from 30 measurements using the PyRosetta output packstat function. The controls were the structures generated using the modified protocol from [13]. For additional comparison the top7 (PDB ID: 1QYS) [22] protein packing stat is measured along with the five structures from the protocol paper by [13].

| Structure | Packing Score | Structure | Packing Score | Structure | Packing Score | Structure | Packing Score |
| --- | --- | --- | --- | --- | --- | --- | --- |
| 1 | 0.610 | 11 | 0.612 | 21 | 0.803 | top7 | 0.479 |
| 2 | 0.657 | 12 | 0.673 | 22 | 0.758 | 2KL8 | 0.447 |
| 3 | 0.647 | 13 | 0.520 | 23 | 0.489 | 2LN3 | 0.630 |
| 4 | 0.632 | 14 | 0.713 | 24 | 0.693 | 2LTA | 0.627 |
| 5 | 0.548 | 15 | 0.698 | 25 | 0.705 | 2LV8 | 0.700 |
| 6 | 0.663 | 16 | 0.660 | Control 1 | 0.564 | 2LVB | 0.635 |
| 7 | 0.655 | 17 | 0.756 | Control 2 | 0.534 |  |  |
| 8 | 0.731 | 18 | 0.788 | Control 3 | 0.568 |  |  |
| 9 | 0.638 | 19 | 0.720 | Control 4 | 0.555 |  |  |
| 10 | 0.734 | 20 | 0.649 | Control 5 | 0.501 |  |  |

**Figure 1.**
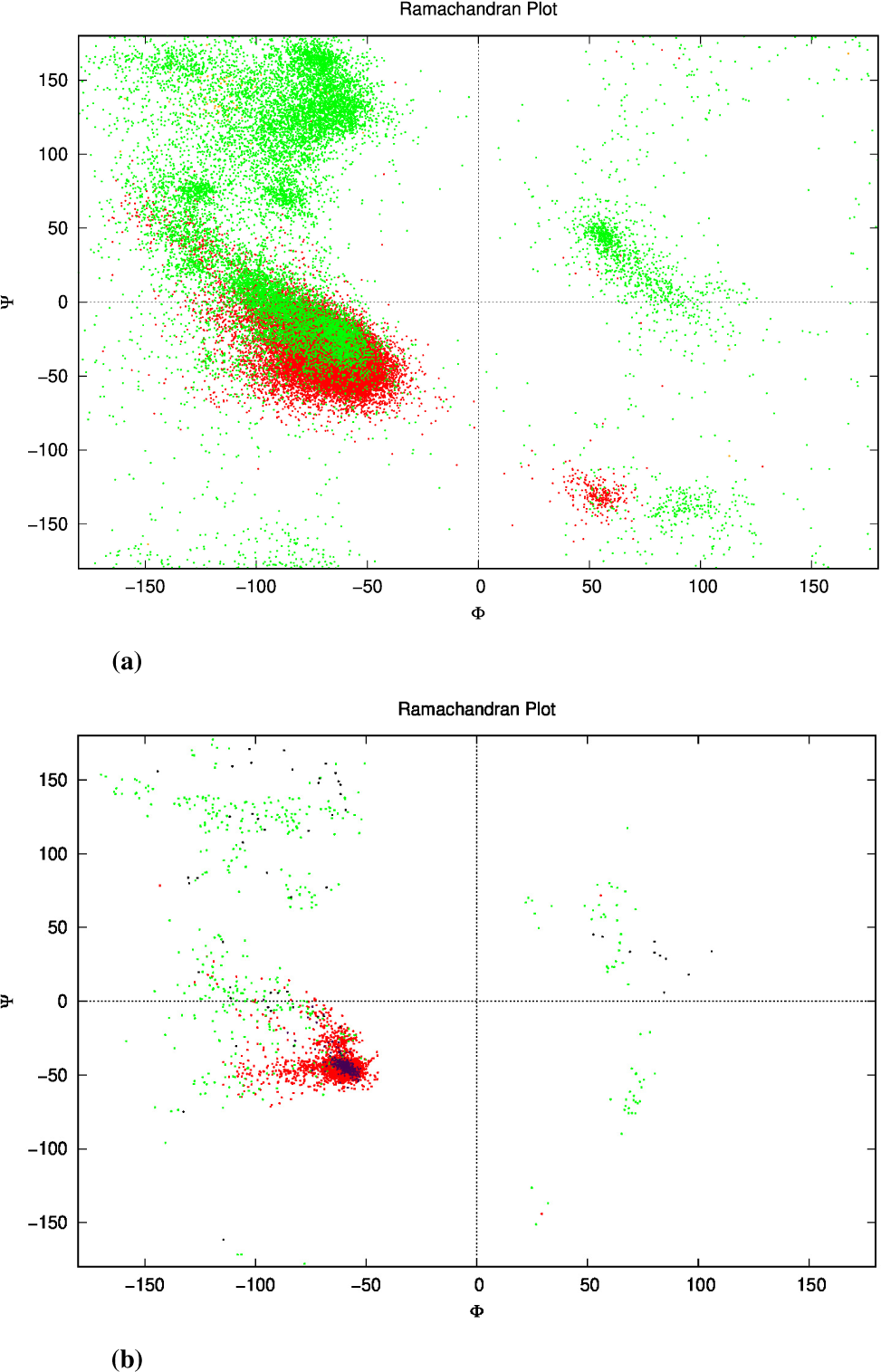
Ramachandran plots. 7A: Ramachandran plot of the dataset showing the *ϕ* and *ψ* angles of each amino acid for each structure. This is the un-augmented data of structures that are only made of helices. Green represents the angles on amino acids in loops, while red represents the angles of amino acids in helices. Some orange can be seen where the DSSP algorithm classified the amino acids as sheets (though there were none). One point to note; the angles here are represented between the range −180° to 180° as is conventional, while in the actual dataset the range was from 0° to 360°. **7B:** The network’s output *ϕ* and *ψ* angles for 25 structures after the relaxation step. The green dots represent the angles of amino acids within loops, and red within helices clustering around the same location as Fig 1A within the fourth quadrant as is desired for an *α*-helix that has ideal angles around (−60°, −45°). These structures culminated to all 25 structures in Fig 4, and had an angle range for the helices (−127.4°<*ϕ*< −44.7°, −71.3°<*ψ*<30.6°) not including the outliers. The purple dots represent the helices in the control structures, and the black dots the loops.

### The neural network

The model in Figure 2 was built using the SenseGen model as a template [2] and consisted of a network constructed from an LSTM layer with 64 nodes, followed by two dense fully connected MLPs with 32 nodes for the first layer and 12 nodes for the second one, both employed a sigmoid activation function:

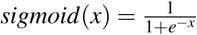

Which was followed by an MDN layer employing an MDN activation function:

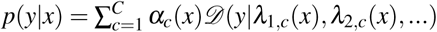

*c*: the index of the corresponding mixture component. *α*: the mixing parameter. 𝒟 : the corresponding distribution to be mixed. *λ* : the parameters of the distribution 𝒟, as we denote 𝒟 to be a Gaussian distribution, *λ*_1_ corresponds to the conditional mean and *λ*_2_ to the conditional standard deviation. The training was done using the Adam optimiser [30], for each parameter *ω* ^*j*^ :

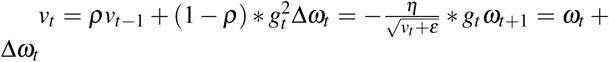

*η*: initial learning rate. *v*_t_: exponential average of squares of gradients. *g*_t_: gradient at time *t* along *ω* ^*j*^. The loss defined as the root mean squared difference between the sequence of inputs and the sequence of predictions:

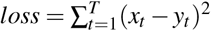

*y*_t_: output. *x*_t_: next step sample *x*_t+1_ = *y*_t_.

**Figure 2.**
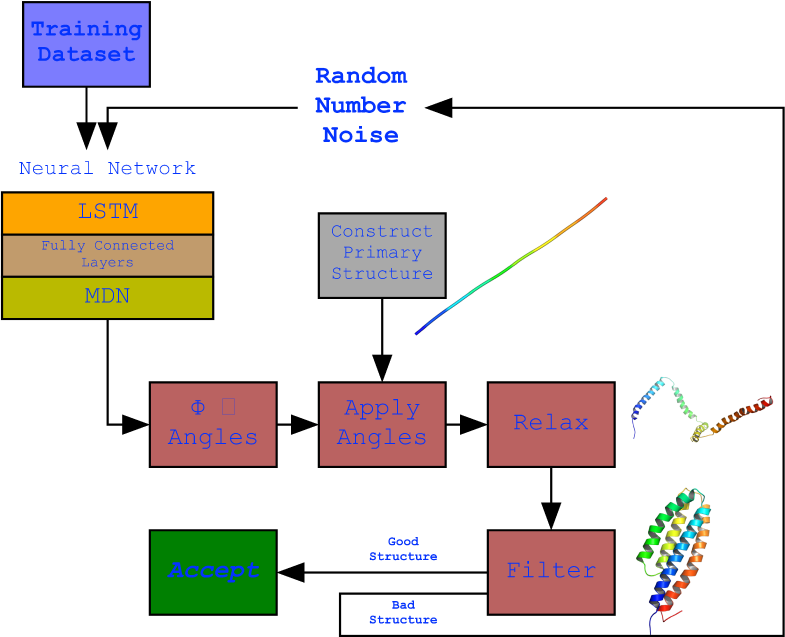
The *de novo* helical protein backbone design protocol. The full protocol showing the structure of the neural network model and its output. The model employing an LSTM network. The network’s output is the generated *ϕ* and *ψ* angle sequences which were applied to a primary structure (a fixed 150 valine length straight structure generated by PyRosetta) that resulted in the development of the secondary structures but not a final compact structure structure due to suboptimal loop structures as a result of their variability in the dataset. To overcome this, the structure was relaxed to bring bring the secondary structure helices together. This did result in more compact structures but was not always ideal, thus a filter was used to filter our non-ideal structures and keep an ideal structure when generated.

The network used random noise as a starting seed, this noise was generated by taking a single randomly distributed number between [0, 1) as the first p redicted v alues, these values were then reshaped to the same shape of the last item of the predicted value resulting in a final shape of (batch size, step number, 1). The network predicted the main parameters of the new value (*µ, σ, π*) several times (according to the number of mixtures value) and selected the single mixture randomly but according to the *π* value. It then predicted the next value according to the normal distribution using the *µ* and *σ* values. It added the final value to the prediction chain and then returned to step 2 until the predefined sequence length was obtained. The initial random number was stripped from the returned sequence. Once the networks were constructed, the dataset was normalised and the training was done using the following parameters: the learning rate was 0.0003 with a batch size of 4 over 18,000 epochs.

### Post-backbone topology generation processing and filtering

The output of the neural network was always a sequence of 150 *ϕ* and 150 *ψ* angle value combination for a structure with 150 amino acids. A straight chain of 150 valines was computationally constructed using PyRosetta and used as a primary structure. Each amino acid in the primary structure had its angles changed according to the *ϕ* and *ψ* angle values, which ended up folding that primary structure resulting in secondary structures of helices and loops between them. The COOH end was trimmed, if it was a loop, until it reached the first amino acid that comprised a helix, thus variable structure sizes where generated. The structure ended up with helices and loops yet still with an open conformation. The generated structure was therefore relaxed using PyRosetta version 4 FastRelax protocol to idealise the helices and compact the structure. Furthermore, not every prediction from the neural network resulted in an ideal structure even after the relax step, therefore we employed a filter to filter out structures we deemed not ideal. The filter discards structures that were less than 80 amino acids, had more residues in loops than in helices, had less than 20% residues making up its core, and had a maximum distance between C*α*1 and any other C*α* greater than 88 Å (the largest value in the dataset). PyRosetta was used [3] since it was easier to integrated the code with the neural network’s python script and combine the Rosetta engine with Biopython [4] and DSSP [12] [18].

### Rosetta *de novo* protein design as a comparison

As a control we used the *de novo* protein design protocol using the Rosetta Script from the [13] paper’s supplementary material. The protocol was modified slightly to accommodate new updates in the Rosetta software suite, but maintained the talaris2014 energy function as in the original paper, and we used the protocol to design helical proteins. These proteins had to pass through several filters including a talaris2014 score of less the -150, a packing threshold of more than 0.50, and a secondary structure threshold of more than 0.90. We attempted to design proteins with 3, 4, and 5 helices to compare the backbone quality our neural network output to the output of the [13] paper.

### Implementation

This setup used the following packages: python 3.6.9, PyRosett 4, Rosetta 3, Tensorflow 1.13.1, BioPython 1.76, and DSSP 3.01 and was run on GNU/Linux Ubuntu 19.10. Further information on running the setup can be found on this GitHub repository which includes an extensive README file. To train the neural network a 3GB GPU and 12GB of RAM is recommended, while to run the trained neural network and generate a backbone structure an Intel i7-2620M 2.70GHz CPU and 3GB of RAM is recommended.

### Operation

As detailed in Figure 2, executing the trained network will generate a set of random numbers that are pushed through the network which (using the weights) will modify the values to become values in accordance with the *ϕ* and *ψ* angle topologies observed in the training dataset. A simple straight chain of 150 valines is then computationally constructed and the *ϕ* and *ψ* angles are applied to each amino acid in order, resulting in the appearance of helical secondary structures. Any tailing loops at the COOH end will be cut out since it interferes with the next step. This structure is then relaxed using the FastRelax protocol which moves the *ϕ* and *ψ* angles randomly in its attempt to find the lowest scoring configuration compacting the structure in the process. A filter is applied to determine whether the final structure is ideal or not, its parameters are detailed in the Methods section. If the structure passes the filter the script exits, otherwise it repeats the whole process.

## Results

The dataset was named PS Helix 500 due to the fact that the features used were the *ϕ* and *ψ* angles, only strictly helical protein structures were used, and each structure was augmented 500 times.

Minimal hyperparameter search was needed and was peformed manually. The neural network was trained on the dataset for 18,000 epochs (further training collapsed the network, i.e. all outputs were exactly the same structure) with a generally sloping down mean loss as shown in Figure 3 indicating that the network got better at generating novel data compaired to the original training data. The network was used to generate a sequence of *ϕ* and *ψ* angles for 25 structures. In other words, random numbers were generated then pushed through the network where they were modified (using the network’s trained weights) to become the *ϕ* and *ψ* angles for 150 residues. This is the angle profile. Using PyRosetta, a simple straight 150-valine chain was constructed and used as a primary structure. The generated *ϕ* and *ψ* angles were applied to this primary structure (each amino acid’s angles were changed according to the angle profile) resulting in a folded structure clearly showing helical secondary structures with loops between them. The last tailing loop at the COOH end was truncated, resulting in structures with variable sizes. The loop angles were within the area of the angles found in the dataset 1, but because they were generated independent of the thermodynamic stability of the structure, the structure did not come together into compact a topology. To push the structure into a thermodynamic energy minima it was relaxed using the PyRosetta FastRelax function which compacted the backbone topology while still having valines as a temporary placeholder sequence. This was repeated 25 times to result in the 25 structures in Figure4. For comparison five control structures were generated using the *de novo* design protocol by the previous paper [13]. Figure 1B shows the Ramachandran plot of the 25 generated structures, where red are the amino acids within helices having angles clustering around the same location as Figure 1A (in the fourth quadrant), as is desired for an *α*-helix, which has ideal angles around (−60°, −45°), our results had an angle range for the helices (−127.4° <*ϕ*< −44.7°, −71.3°<*ψ*<30.6°) not including the outliers, the five control structures show angles within the same region (purple for helices and black for loops). The structure generation setup was not perfect at achieving an ideal structure every time, so a filter was deployed to filter out suboptimal structures by choosing a structure that had more residues within helices than within loops, was not smaller than 80 residues, and had more than 20% residues comprising its core. Due to the random nature of the structure generation the efficiency of the network is variable, while generating the 25 structure in 4 the fasted structure was generated after just 4 failed attempts, while the slowest structure was generated after 3834 failed attampts giving a success rate for the network between 25.0% at its best and 0.025% at its worst, and this took between ∼1 minute and ∼6 hours to generate a single structure with the desired characteristics utilising just 1 core on a 2011 Mac-Book Pro with with 4 core Intel i7-2620M 2.70GHz CPU, 3GB RAM, and 120GB SSD. For comparison the protocol by [13] took ∼60 minutes for each of the control structures on the same machine. The protocol is summarised in Fig 2, and the results are compiled in Fig 4 showing all 25 structures and the 5 controls.

**Figure 3.**
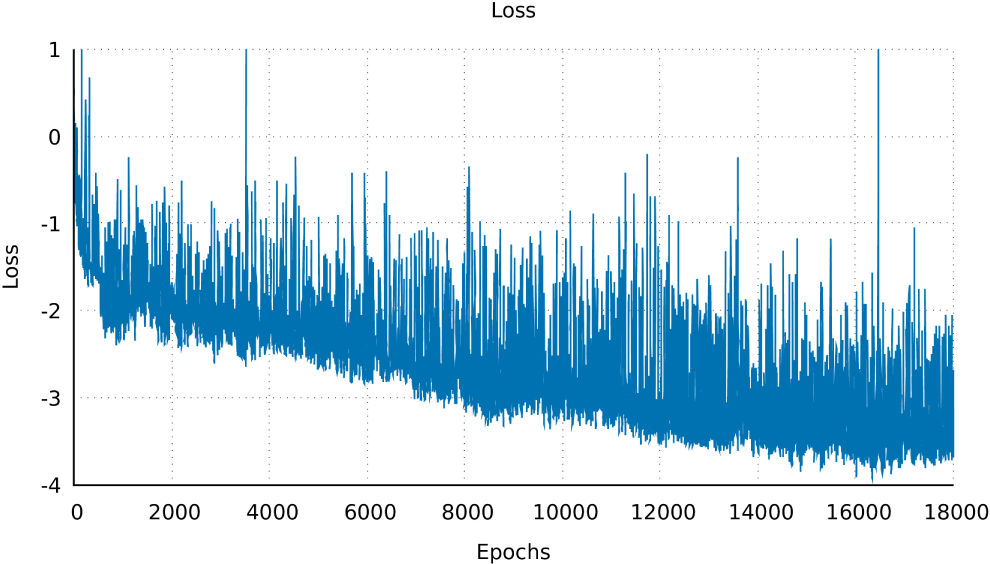
The training loss. The mean loss of the whole network over epoch, for 18,000 epochs, showing a general downward trend. This indicated that in subsequent epochs the network got better at generating structures.

**Figure 4.**
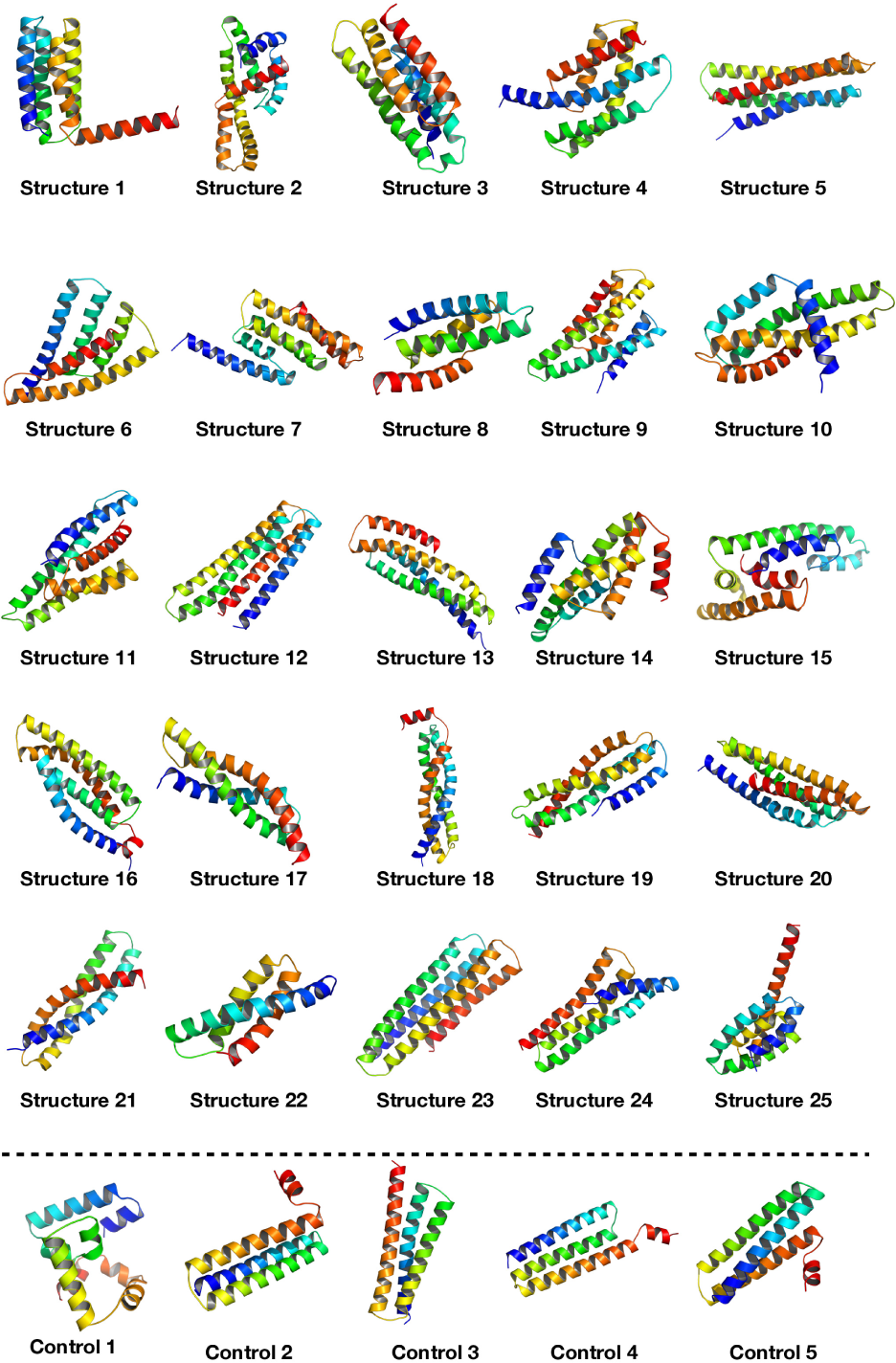
The designed structures. This figure shows all 25 structures that were generated using the neural network, displayed using PyMOL [21]. It can be seen that all structures have compact helical structures of variable topologies. The 5 control structures at the bottom were generated using the protocol from the [13] paper showing better helices but similar compactness.

These 25 structures had on average 84.7% of their amino acids comprising their helices, along with and an average of 29.9% of their amino acids comprising their cores, 2 shows the Rosetta packing statistic for all 25 structures, along with the five controls, the top7 protein, and the five structures from the [13] paper all showing similar packing values.

The LSTM network was not the only network that was tested, an MLP-GAN [32] was also tested on the same dataset, hyperparameter search was performed through a grid search then a random search to finally reach the following hyper-parameters for the generator network: 4 layers with the following nodes 256, 512, 1024, and batch normalization with 0.15 momentum between the layers, and an output node with the shape of the dataset (150, 2). This network employed the ReLU activation for the hidden layers and the TanH activation function for the output layer with the binary crossentropy loss and the Adam optimiser with a 0.0004 learning rate. The discriminator network had the following hyperparameters: 3 layers with the following nodes 512, 256, and 1 node for the output, and 25% dropout rate between the layers. This network employed the ReLU activation for the hidden layers and the sigmoid activation function for the output layer with the binary crossentropy loss and the Adam optimiser with a 0.002 learning rate. The architecture was trained for 100,000 epochs with a mini batch size of 512 and the results were not as good as the LSTM Fig 5; generating structures that did have helices and loops but much more sparse and kinked. Thus the LSTM is the preferred architecture for this particular dataset.

**Figure 5.**
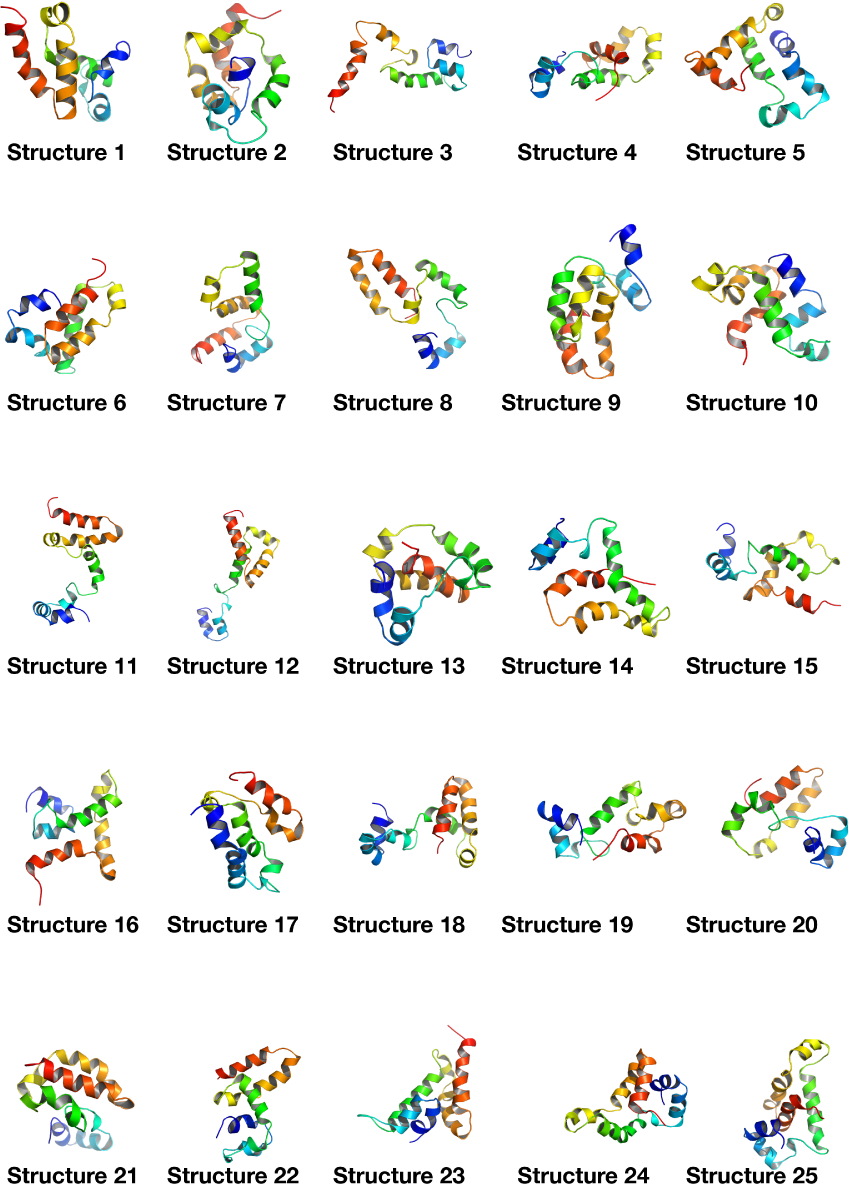
Results from an MLP-GAN. This figure shows 25 structures generated from an MLP-GAN, it can be inferred that these structures are not as good as the ones generated from an LSTM since they are much more fragmented; with smaller kinked helices and numerous loops between them.

## Discussion

In this paper we outlined how a neural network architecture can design a compact helical protein backbone. We attempted to test our network to generate structures that included sheets, but that failed, mainly due to the wide variation in the loop angles that did not compact a structure to bring the strands together, which sheets rely on more to develop compared to helices.

We demonstrated that the *ϕ* and *ψ* angles were adequate features to design a protein backbone topology only without a sequence. Although we understand the distribution of angles in the Ramachandran plot (the distribution of helix and sheet angles) the neural network’s power comes in the form of combining several dimensional spaces to take a better decision. Given its observation of natural proteins, it can calculate the ideal combination of angles that results in deciding the number and lengths of helices, and the loops between them, that will still result in a compactly folded protein backbone.

The reason we concentrated our efforts on generating a backbone only is because once a backbone is developed it can be sequence-designed using other protocols, such as RosettaDesign[24] [25] or the protocol from [20].

Though our network had a wide variation of success rates, from high to low, that was due to the random nature of the setup, which was our target to begin with (to randomly generate backbone topologies rather than directly design a specific pre-determined topology). Generating multiple structures and auto filtering the suboptimal ones provided an adequate setup, this achieved our goal of *de novo* helical protein backbone design within a reasonable time (1-6 hours) on readily available machines.

As a control we used a slightly modified *de novo* design protocol from [13], which also performs backbone design (resulting in an all-valine backbone topology) followed by sequence design of that backbone. It has numerous advantages such as generating better helices, but the user must still predetermine the topology to be generated (decide the number of helices, their lengths and locations, and the lengths of the loops between them), while this neural network automatically takes that decision and randomly generates different backbones, which can be very useful for database generation (see below).

The neural network is available at this GitHub repository which includes an extensive README file, a video that explains how to use the script, as well as the dataset used in this paper and the weights files generated from training the neural network.

For future work we are currently working on an improved model that uses further dataset dimensions and features that will allow the design of sheets.

## Use Cases

There are many benefits to generating numerous random protein structures computationally. One benefit can be in computational vaccine development, where a large diversity of protein structures as scaffolds are required for a successful graft of a desired motif[27] [28] [29]. Using this setup, combined with sequence design, a database of protein structures can be generated that provides more variety of scaffolds than what is available in the protein databank, especially that this neural network is designed to produce compact backbones between 80 and 150 amino acids, which is within the range of effective computational forward folding simulations such as AbinitioRelax structure prediction[26].

## Author contributions

Corresponding author: Sari Sabban Author roles Sari Sabban: Conceptualization, Data Curation, Formal Analysis, Investigation, Methodology, Project Administration, Resources, Software, Supervision, Validation, Visualization, Writing – Original Draft Preparation. Author roles Mikhail Markovsky: Investigation, Methodology, Software, Writing – Review & Editing.

## Acknowledgements

The corresponding author would like to thank the High Performance Computing Center at King Abdulaziz University for making available the Aziz high performance computer where the corresponding author was able to augment the dataset and perform the Abinitio folding simulations.

